# Chromosome-level genome assembly of the Olympia oyster (*Ostrea lurida*), the only oyster species native to the West Coast of North America

**DOI:** 10.64898/2026.09.23.753890

**Authors:** Bernarda Calla, Sheina Sim, Mark S. Yeats, Kevin M. Johnson, Karla Chavez-Congrains, Angela N. Kauwe, Louis V. Plough, Scott M. Geib

## Abstract

The Olympia oyster (*Ostrea lurida*) was once a major food source and a key component of coastal marine ecosystems and culture on the US West Coast. Overfishing and habitat contamination in the early to mid-20th century severely depleted populations of Olympia oysters, and the species is currently considered a species of concern. Although multiple restoration efforts are underway, progress understanding the species biology and recovery potential has been limited by the historical lack of genomic resources. Here, we report a chromosome-level assembly of the Olympia oyster. The assembly was built using Oxford Nanopore reads and Hi-C to obtain a highly contiguous 1.03 Gb-assembly consisting of 10 chromosomes. RNA from gills, mantle, adductor muscle, and a mix of brooding eggs and larvae were also sequenced as 150 nt long paired-end reads which were used to guide the genome annotation. Over 52 K genes were predicted in the genome, of which 34K were protein-coding. A BUSCO analysis on the assembly showed 98.9% of the mollusca_odb12 gene set were complete (with 2.1% duplicated, and 0.8% missing). The short RNA reads were also used to identify tissue-level gene expression signatures in adductor muscles, egg and brooding larvae, and mantle. Development-related genes were highly expressed in eggs and brooding larvae whereas many xenobiotic metabolism genes were present in mantle tissues, highlighting mantle’s role in the interaction of the oyster with external environments. This new genomic resource will greatly advance our understanding of this keystone mollusk and provide a critical tool for guiding restoration efforts and the development of sustainable commercial production.

## Introduction

The Olympia oyster (*Ostrea lurida* Carpenter, 1864) is the only native oyster species on the North American West Coast. This small oyster, averaging only five centimeters in length when fully mature, could once be found covering entire subtidal aquatic beds along the coasts of Washington, Oregon, Alaska, California, and Baja California in Mexico (Steele, 1957). By the end of the 19^th^ century, Olympia oysters became nearly extinct owing to overharvesting and habitat alteration (Zu Ermgassen et al., 2012) (Figure 1).

**Figure 1.**
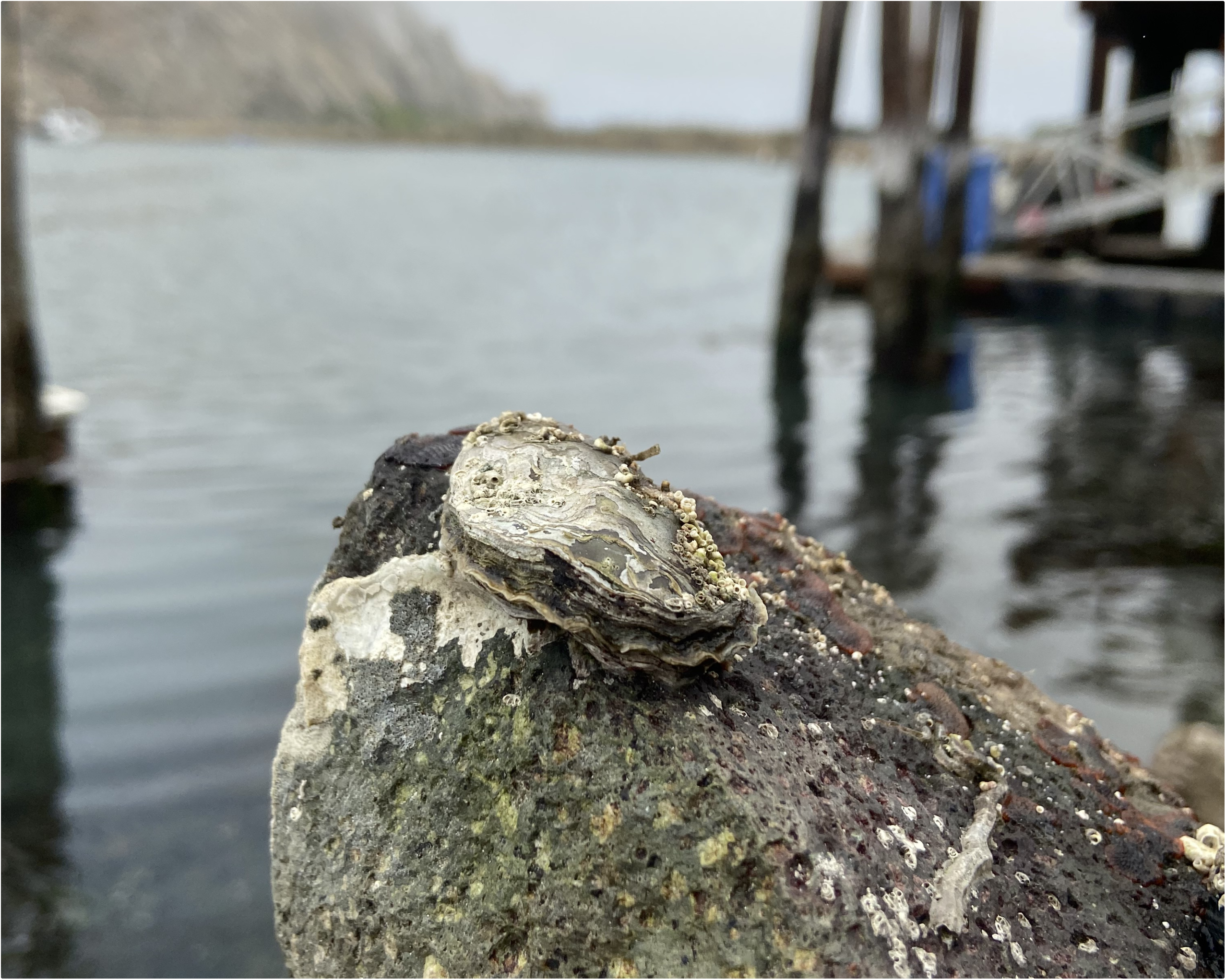
Adult Olympia oyster (*Ostrea lurida*) (Photo: Kevin M. Johnson)

Olympia oysters were an integral part of local ecosystems: besides being a source of food, they were important elements of the estuarine landscape, providing ecosystem services that include water filtration, coastline protection through reef stabilization, and habitat for other species (Blake & Ermgassen, 2015; Pritchard et al., 2015; White et al., 2009). But these oysters were also part of an economic and cultural network, closely tied with Indigenous peoples’ traditions and rituals, and sustainably harvested and traded as a valuable commodity (Echtle, 2014; Grier et al., 2017; Luby & Gruber, 1999; Reeder-Myers et al., 2022). In the mid to late 19^th^ century, Indigenous and non-native settlers in the West Coast entered agreements that granted ownership and use rights to tidelands where Olympia oysters naturally occurred, provided that these stablished oysters populations were preserved (Echtle, 2014; Steele, 1957). Soon after, cultivation techniques were developed to scale up production, greatly increasing yields, distribution, and consumption. However, expanding human activities, including natural resources overexploitation, the growth of polluting industries, and the introduction of foreign pest species along with the arrival of the competing and easier to cultivate Pacific oyster (*Crassotrea gigas*) in the early 1900, rapidly led to the decline of the Olympia oyster industry, and pushed the species to the brink of extinction (Steele, 1957, 1964).

Currently, Olympia oysters are found mainly in small, isolated, and spatially dispersed reefs within coastal estuaries (e.g. Blake & Ermgassen, 2015). Most existing populations are closely tied to restoration activities that began in the early 2000s and are concentrated in Washington, California, and, to a lesser extent, Oregon (Barber et al., 2015; Blake & Bradbury, 2012; Ridlon, Marks, et al., 2021; Wasson et al., 2020). Although the focus of these efforts is largely to rebuild natural populations and restore estuarine habitats, some work has examined the species’ potential for aquaculture production and consumption (e.g., Baker, 1995; Ridlon, Wasson, et al., 2021).

Oysters in the *Ostrea* genus (flat oysters) are sequential hermaphrodites, spawning first as males and then alternating between sexes in subsequent spawns (Coe, 1932). Unlike oysters in other genera, such as *Crassostrea* or *Magallana* (cupped oysters), *Ostrea spp.* fertilize their eggs within the female oyster and brood those fertilized eggs inside their mantle cavity up until the veliger larval stage when larvae are released and can spend weeks in the plankton before settlement. After settlement, *O. lurida* grows relatively slowly compared to cupped oysters, typically reaching only 5–8 cm in maximum shell length over several years, depending on environmental conditions (Pritchard et al., 2015).

Although slow growth, coupled with shifting habitat conditions and disease pressure, seems to greatly constrain restoration efforts (Friedman et al., 2005; Trimble et al., 2009), other life history traits of Olympia oysters appear to be advantageous. It has been proposed that early exposure of *Ostrea* spp. larvae to fluctuating brood-chamber conditions—especially reduced pH—may increase larval resistance to ocean acidification (OA). Gray et al. (2026; 2019) suggested that this chemically variable environment pre-conditions larvae, giving *Ostrea* spp. greater acclimatory potential under future OA compared with cupped oysters and other broadcast-spawning shellfish. Continued research on *Ostrea spp.* physiology, genetics, and genomics is needed to identify the mechanisms behind this increased resilience and the ways it may impact population persistence as environmental conditions in coastal environments continue to change.

Despite the great ecological and cultural significance of Olympia oysters, there is a striking dearth of genomic resources available for this species, especially when compared to other species undergoing significant restoration efforts like the Eastern oyster (*Crassostrea virginica)* in the U.S. (e.g. Guo et al., 2023; Puritz et al., 2024), or the Olympia oyster congener in Europe: the European flat oyster (*Ostrea edulis*) (e.g. Bean et al., 2022; Boutet et al., 2022). Although a single study has characterized the population genetic structure of Olympia oysters across their U.S. West Coast range (Silliman, 2019), the functional genomic context underlying this variation remains unresolved in the absence of a reference genome. Here, we make available the first chromosome-level assembly of the Olympia oyster. This genome resource will accelerate research on the species by enabling detailed assessments of genetic diversity in existing populations, quantifying inbreeding levels, and improving long-term monitoring efforts—ultimately supporting restoration and conservation initiatives.

## Methods

### Specimen collection and dissections

Year-class 2022 adult and year-class 2024 juvenile Olympia oysters were received through the Puget Sound Restoration Fund (PSRF) from the NOAA NW Fisheries Research Station in Port Orchard, WA on June 6, 2024. Prior to shipping, oysters were kept in hatchery conditions at 16°C and ∼28-29 ppt. Oysters were sourced from Clam Bay, Central Puget Sound, WA, and a Shellfish Mariculture Transport Permit (#24-008) was approved through the Oregon Department of Fish and Wildlife on May 28, 2024. All animals were certified as healthy prior to procurement (AquaTechnics AQ24-57). Adult oysters (n=7) were 2 years old, and at a size range of 20-30 mm in length. Juvenile oysters (n= 14) were 4 months old, and 10-15 mm in length. Upon arrival, all animals were placed in a quarantine trough in the Newport Aquaculture Lab (NAL) Disease Wing at Oregon State University’s Hatfield Marine Science Center (HMSC). Oysters remained in quarantine from June 6, 2024, until dissections on June 24, 2024. Water temperatures were checked daily in the quarantine system. Temperature remained at a constant 22 °C for the first week and was lowered to an average temperature of 19.63 ± 0.48 °C until dissections. Salinity remained at ∼30-32 ppt over the quarantine period. On June 24, 2024, animals were dissected one at a time in petri dishes over wet ice. Instruments used for dissections (shucking knives, micro scissors, forceps, and scalpels) were rinsed with DI water before and after a 10% bleach rinse between each animal. Whole bodies were collected for the juvenile animals, while adductor, eggs and brooding larvae, ctenidia (gills), and mantle tissues were collected from adults. Only three adult oysters had gonads that were mature for sexing. All three mature adults were female. For two mature females, pooled brooding clutches were collected consisting of recently fertilized eggs and veliger larvae found within each female upon shucking. All tissue collections were placed in nuclease-free 1.5 mL microcentrifuge tubes prior to flash freezing with liquid nitrogen.

All tissue collections were stored at – 80 °C immediately after flash freezing. On July 9, 2024, whole body juveniles (n=4), adductor tissue (n=2), mantle tissue (n=2), ctenidia tissue (n=1), and both brooding clutches immature brood collections were shipped overnight on dry ice to the USDA-ARS Tropical Pest Genetics and Molecular Biology Research Unit in Hilo, Hawaii.

### Nucleic acid extraction, library, preparation, and sequencing

High molecular weight (HMW) DNA extraction was performed using a salt-out protocol derived from Miller, et al., (Miller et al., 1988). Following nucleic acid isolation, the HMW genomic DNA was size-selected using the Circulomics Short Read Eliminator XS or XL (PacBio) kit to remove short fragments. A native DNA library was prepared from 900–1200 ng of DNA using the Oxford Nanopore Technologies Ligation Sequencing Kit V14 (SQK-LSK114) according to the manufacturer’s instructions and sequenced on a PromethION 24. Sequencing was performed on R10.4.1 (FLO-PRO114M) flow cells with live base calling implemented through MinKNOW using lDorado and Super Accurate Basecalling (SUP). Resulting fastq files were concatenated and used for downstream genome assembly.

Concurrent to the WGS library preparation, Hi-C library preparation was performed using tissue from one juvenile female individual. This tissue was cross-linked, digested using the enzymes DdeI and DpnII, and proximity ligated. After proximity ligation and nucleic acid purification, the sample was size-selected to enrich for DNA fragments 200-600 bp. Following sample purification using SPRI-beads, library preparation was performed using the NEB Next Ultra II DNA Library Prep Kit. The resulting Hi-C library was sequenced on one 2 x 150 Cloudbreak FS Sequencing Kit on an Element Biosciences Aviti system to approximately 400,000 proximity ligations.

Total RNA was extracted from one juvenile female through homogenization of snap frozen tissue in Tri Reagents and then extraction using the Zymo Direct-zol Magbead Total RNA kit (Zymo Research, Irvine, CA) on a Kingfisher Flex 96 system (ThermoFisher, Waltham, MA). From total RNA, a poly(A) cDNA library was generated using the NEB Ultra II RNA Library Prep Kit (New England Biolabs, Ipswich, MA) using oligo(dT) selection. The resulting library was sequenced on one 2 x 150 Cloudbreak FS Sequencing Kit on an Element Biosciences Aviti system.

### Genome assembly and annotation

The Oxford Nanopore reads were concatenated into a single file and reads were assembled into contigs using HiFiASM (v.0.24.0-r702) (Cheng et al., 2021; Cheng et al., 2022) with implementation of the –ont flag for standard FASTQ input. The resulting primary assembly, alternate assembly, and error corrected reads produced by HiFiASM were supplied to PurgeDups (Guan et al., 2020) to remove duplicate contigs from the primary assembly. The duplicate purged contig assembly served as the reference for the Hi-C scaffolding pipeline which implemented BWAmem 2 (v.2.2.1) (Vasimuddin et al., 2019) for mapping Hi-C reads to the reference, Picard (v.3.2.0) (Picard2019toolkit, 2019 https://github.com/broadinstitute/picard) for removing PCR duplicates, and YAHS (Zhou et al., 2023) for creating a Hi-C contact map. The resulting Hi-C contacts were visualized and manually curated using Juicebox (v.2.15) (Durand et al., 2016).

Taxonomic assignments for all records in the assembly were identified using NCBI Foreign Contamination Screen (FCS-gx) (https://github.com/ncbi/fcs-gx) (Astashyn et al., 2024) and non-mollusc contigs were separated from the assembly using FastaParser (https://github.com/sheinasim-USDA/FastaParser). Visualization of the assembly was performed using Blobtools2 (Challis et al., 2020). This included estimation of completeness of the genome using BUSCO (v.6.0.0) (Manni et al., 2021; Tegenfeldt et al., 2025) with the mollusca_odb12 gene set.

Genome annotation of the primary *O. lurida* assembly was done with the Eukaryotic Genome Annotation Pipeline - External (EGAPx) (https://github.com/ncbi/egapx) v0.5.2 and short-read RNA-seq data as evidence. Briefly, EGAPx aligned the tissue-specific transcriptome reads reference using STAR v0.15 and used Miniprot v2.7.11 to align the Mollusca protein set to the reference. The pipeline used gnomon (https://www.ncbi.nlm.nih.gov/refseq/annotation_euk/gnomon/) for gene prediction using the resulting alignments and additionally made *ab initio* gene predictions based on HMM. Functional annotations to the final structural annotation set were based on orthology and model type and quality. As with the genome, the resulting *O. lurida* annotated gene set was assessed for completeness using the BUSCO mollusca_odb12 gene but with the -prot option. Using the annotated gene set, analysis for synteny with the chromosome assembly of *Ostrea edulis* (NCBI RefSeq Accession: GCF_947568905.1) was performed using Orthofinder v2.5.5 (Emms & Kelly, 2019). Collinear blocks of shared single-copy orthologs were visualized using the R package Genespace (v1.2.3) (Lovell et al., 2022) in R v4.6.1 (R Core Team, 2026).

### Structural heterozygosity

The single file Oxford Nanopore reads were realigned to the assembly using minimap2 v2.31-r1302 (Li, 2018) with the map-ont preset, and alignments were coordinate sorted and indexed using SAMtools v1.24. (Danecek et al., 2021). Separate alignment sets were generated for coverage and breakpoint analyses. Copy-depth calculations used primary alignments with mapping quality ≥20, excluding unmapped, secondary, supplementary, quality-control-failed and duplicate records, whereas structural-variant calling retained supplementary alignments but excluded unmapped, secondary and quality-control-failed records. Repetitive regions were identified *de novo* using Red (REpeat Detector) v5-22-2015 (Girgis, 2015) and excluded together with assembly gaps, the terminal 10 kb of each contig, mitochondrial sequence, and contigs shorter than 100 kb. Coverage was calculated using mosdepth v0.3.14 (Pedersen & Quinlan, 2018) across callable portions of fixed, non-overlapping 10-kb windows containing ≥50% callable sequence. Baseline coverage was estimated as the robust median depth across chromosome-scale scaffolds after excluding values more than three median absolute deviations from the median.

Gene dosage was calculated preferentially from exons with ≥80% callable sequence and at least 100 callable base pairs. Normalized depth was calculated relative to the chromosome-scale baseline, and diploid copy number was estimated as twice the normalized depth. Structural variants were identified using Sniffles2 v2.8.0 (Paulin et al., 2026) with minimum event length and read-support thresholds of 50 bp and 10 reads, respectively, and DELLY v2.6.0 (Rausch et al., 2012) in Oxford Nanopore long-read mode. DELLY was also used to identify read-depth segments, with stringent depth calls requiring PASS, quality >200, uniqueness ≥0.50 and low-complexity fraction ≤0.20. Deletion- and duplication-direction calls were consolidated when their breakpoints occurred within 500 bp or their intervals exhibited ≥50% reciprocal overlap. Candidate events were classified according to breakpoint-caller agreement, concordant depth support, callable fraction, proximity to contig ends and assembly gaps, repeat content and gene-depth direction.

### Tissue-level differential expression

The short-reads RNA-seq libraries for adductor, mantle, and brooding tissues (2 of each) were mapped to the newly assembled genome using Salmon v.1.11.4 (Patro et al., 2017) at the gene level using a “gentrome” and “decoy” sequences to prevent alignment to non-coding regions of the genome. The Salmon output with mapping information for each library was imported into an R object using the tximport package v.1.26.1 (Soneson et al., 2016). The tximport object was then used to run DESeq2 v. 1.38.3 (Love et al., 2014) with the *DESeqDataSetFromTximport* function which directly deals with rounding and uses the length matrix to account for gene length offsets. Differential expression across tissues was calculated using the Likelihood Ratio Test from DEseq2. Genes that were significantly differentially expressed (DEG) between tissues were filtered with an adjusted FDR-corrected p-value cutoff < 0.05, and fold-change > 2. Clustering of DEGs was done with pheatmap v.1.0.12 (Kolde, 2025) with defaults (clustering method: complete, and Euclidean distances). The number of clusters (K) was pre-defined by first running hclust (R Core Team, 2026) and plotting a dendrogram to visualize the tree structure.

## Results and Discussion

### Genome assembly and gene annotation

The final scaffolded assembly had a total length of 1.03 Gb and was comprised of 10 scaffolds designated as chromosomes. These chromosomes represented 0.995 Gb (96.7% of the total assembly). Contigs and scaffolds that could not be placed in a chromosomal context represented a total of 0.0337 Gb (3.3% of the total assembly) (Table 1). The NCBI Foreign Contamination Screen designated a total of 220 contigs in the assembly as derived from prokaryotic viruses, one contig from a nematode, and one contig from a bacterium. These records were removed from the assembly prior to NCBI GenBank submission (Figure 2).

**Figure 2.**
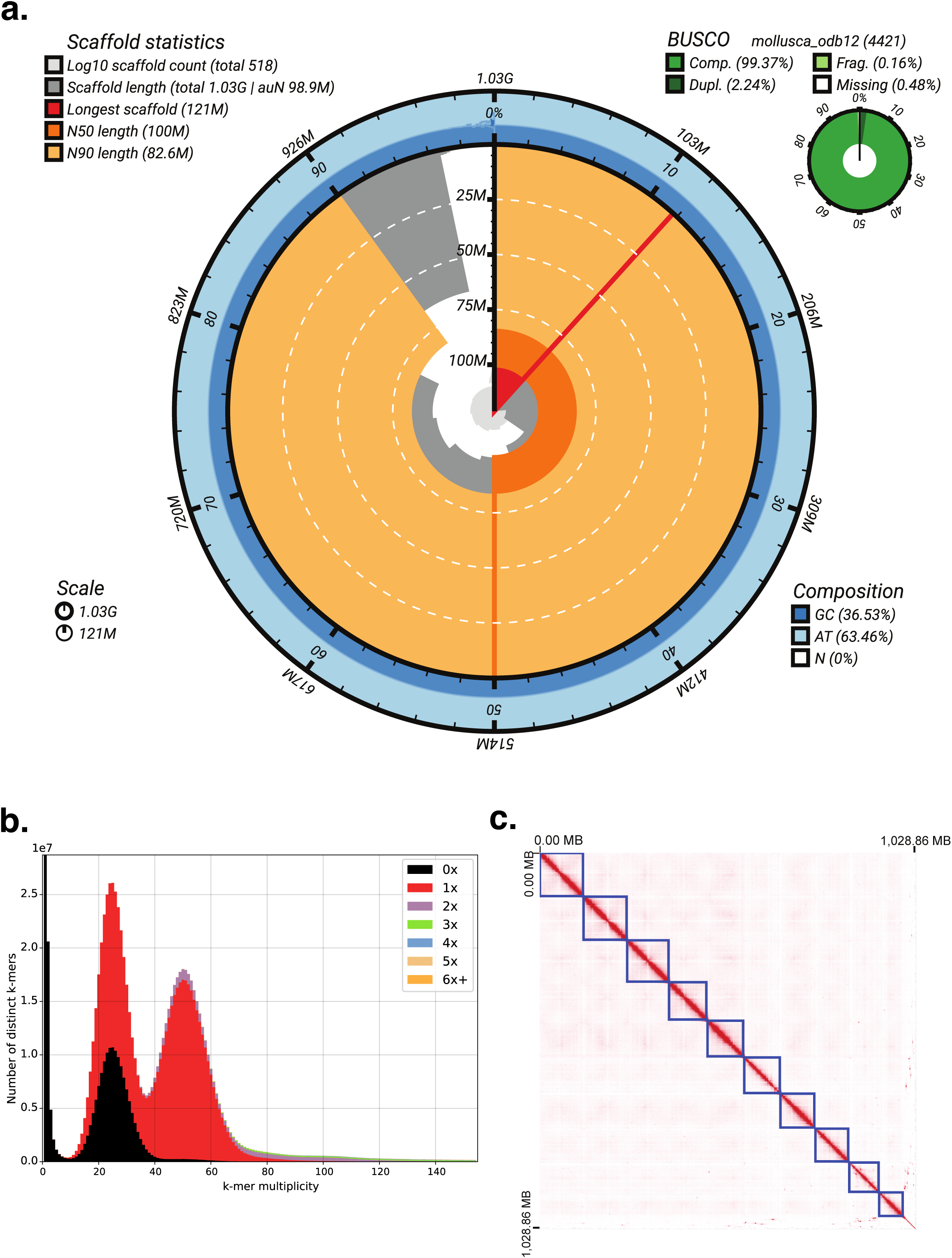
Genome assembly result statistics. **a.** Snail plot visualization of assembly continuity, including BUSCO results with the mollusca_odb12 orthologs database; **b.** K-mer distribution analyses obtained with GenomeScope (Vurture et al., 2017) showing homozygous and heterozygous peaks at ∼50x and 25x coverage; **c.** Hi-C contact map showing the grouping of the ten chromosomes.

**Table 1.**
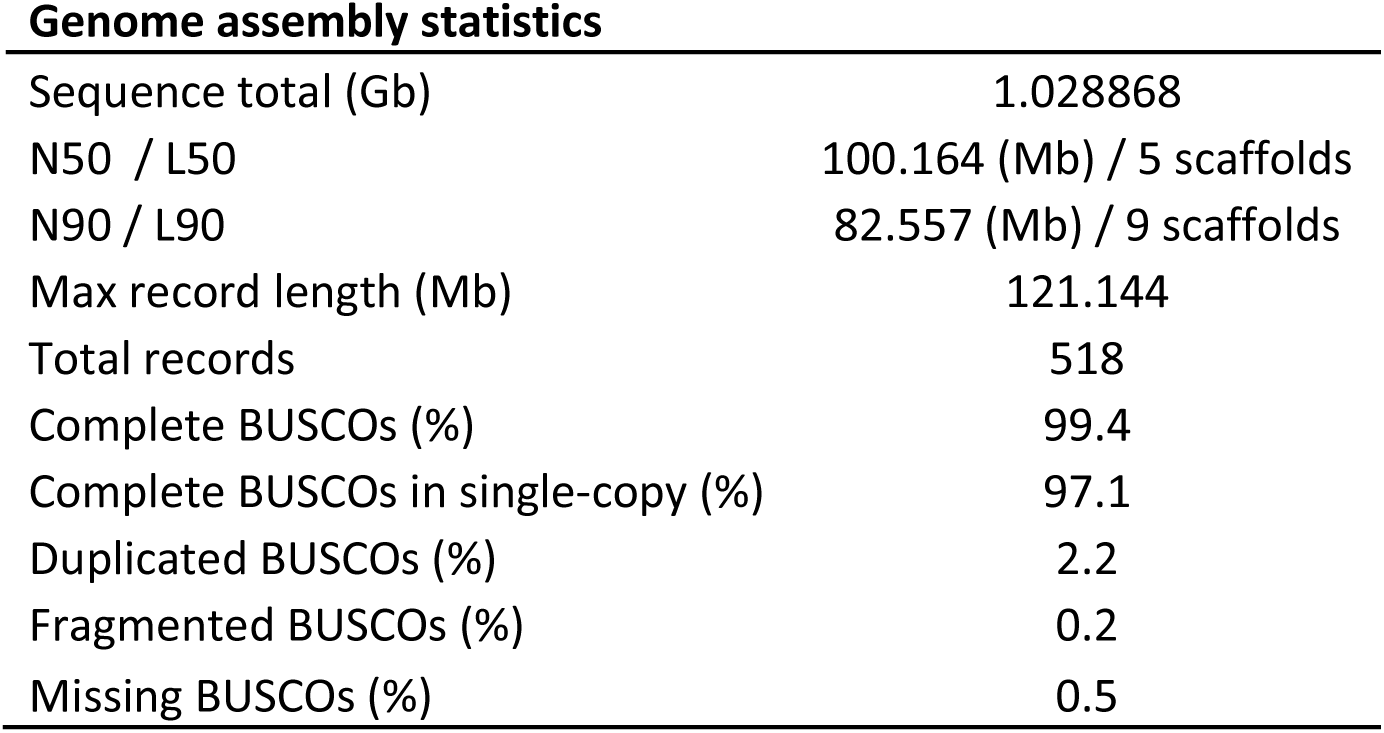
Genome assembly statistics for *Ostrea lurida*.

| **Genome assembly statistics** |  |
| --- | --- |
| Sequence total (Gb) | 1.028868 |
| N50 / L50 | 100.164 (Mb) / 5 scaffolds |
| N90 / L90 | 82.557 (Mb) / 9 scaffolds |
| Max record length (Mb) | 121.144 |
| Total records | 518 |
| Complete BUSCOs (%) | 99.4 |
| Complete BUSCOs in single-copy (%) | 97.1 |
| Duplicated BUSCOs (%) | 2.2 |
| Fragmented BUSCOs (%) | 0.2 |
| Missing BUSCOs (%) | 0.5 |

Gene annotation revealed 52,714 genes, 33,912 of which were protein coding (Table 2). Of the 48,801 annotated mRNA, 42,842 were fully supported (87.8%) and all the 48,801 annotated CDS were fully supported. Assessment of completeness using BUSCO showed 98.9% of the mollusca_odb12 gene set were complete, with 2.1% duplicated, and 0.8% missing. All Mollusca BUSCOs designated as complete were found within the 10 assembled chromosomes. Analysis for synteny using single copy orthologs revealed a highly conserved 1:1 chromosomal orthology between *O. lurida* and *O. edulis* with a high rate of intra-chromosomal translocations and inversions. There were no collinear blocks of three or more genes that were translocated between chromosomes (Figure 3).

**Figure 3.**
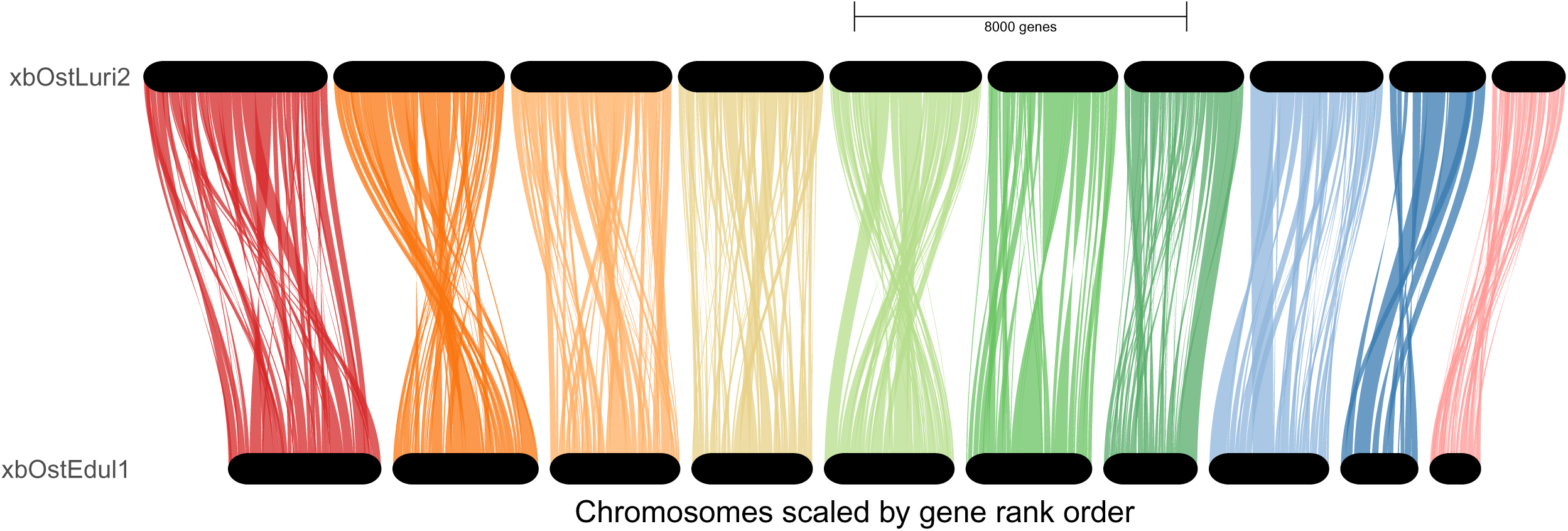
Synteny plot showing conserved chromosomal macrosynteny with extensive intra-chromosomal rearrangements between the newly assembled *Ostrea lurida* genome and the *Ostrea edulis* (European flat oyster; NCBI RefSeq accession: GCF_947568905.1-RS_2023_05).

**Table 2.** Summary of genome annotation results.

| Feature | Count |
| --- | --- |
| **genes** | **52714** |
| genes (non-transcribed pseudo) | 16277 |
| genes (non-coding) | 2525 |
| genes (protein coding) | 33912 |
| **mRNAs** | **48801** |
| **non-coding RNAs** | **3498** |
| **pseudo transcripts** | **16290** |
| **CDSs** | **48801** |

### Structural heterozygosity

Alignment of the Oxford Nanopore reads to the final scaffolded assembly produced a mapping rate of 95.77%. Median coverage across the 6,609 chromosome-scale 10-kb windows was 61.0×. After excluding repetitive sequence, assembly gaps, contig ends, mitochondrial sequence and short contigs, 447.3 Mb, or approximately 43.4% of the assembly, remained callable. Coverage was not strongly associated with GC content within retained windows (Pearson’s r=0.002).

After excluding events shorter than 50 bp, 108,596 read–assembly structural discordances were identified, comprising 106,537 deletion-direction and 2,059 duplication-direction events. Most events (93,680; 86.3%) were classified as artifact-prone, principally because they occurred in regions with insufficient callable sequence. Among the remaining events, 4,336 received moderate provisional support, 10,579 were supported by a single evidence class, and one received high provisional support. 73 high or moderate-provisional events overlapped 83 genes with copy-depth changes in the corresponding direction: 55 intersected annotated exons and 46 completely encompassed at least one gene. The strongest candidate was a heterozygous 6.2-kb deletion on Chromosome 1 supported by identical Sniffles and DELLY breakpoints, a Sniffles allele fraction of 0.525, and a DELLY depth-based copy-number estimate of 1. The deletion encompassed an uncharacterized three-exon protein-coding gene and an adjacent pseudogene (Supplementary table 1).

### Tissue-specific expression

RNA-seq short reads from three different tissues mapped to 52,223 of the predicted genes. From this total, 45% of genes had mapped reads in all tissues, and 1,147 genes were unique to adductor muscle (exactly zero counts across all other tissues, and > 5 raw reads in all replications of that tissue); 218 were unique to brooding clutches, and 3,353 were unique to mantle tissue (Figure 4.A). Using a p-value <0.05 and fold-change > 2 as cutoff, we identified 1,215 genes with differential expression (DE) between the adductor, brooding tissues, and mantle tissues. (Figure 4.B).

**Figure 4.**
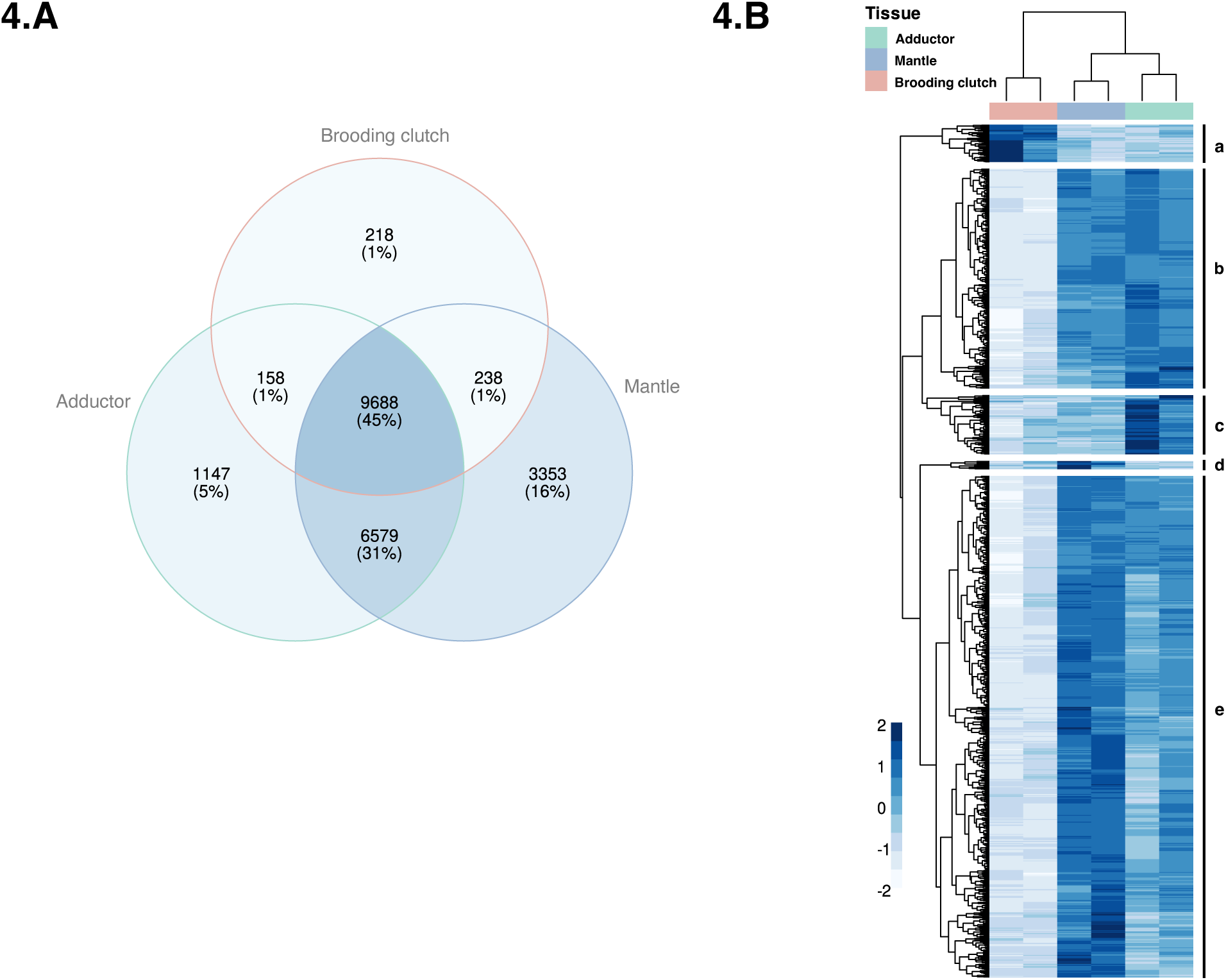
Tissue-level gene expression. **a.** Venn-diagram showing the total number of genes found on three individual *Ostrea lurida* tissues and genes shared between tissues **b.** Heat map showing the hierarchical clustering (K=5) of genes that are differentially expressed between the three tissues; (differential expression cutoff between pairs of tissues: fold-change > 2 and FDR p-value < 0.05).

At the time of dissection of gonadal tissue, it was determined that the adult females were brooding eggs and larvae. Hierarchical clustering of DE genes (Figure 4.B) clearly discriminated genes with higher expression in those brooding clutches (55 genes, 23% unknown function; 3% unannotated). This cluster consisted of genes associated with cell division, development, early embryogenesis, and growth (cluster a). A second cluster (cluster b) grouped genes that were higher in expression in mantle and adductor tissues compared to the brooding clutches tissue (323 genes; 56% unknown function; 4% unannotated). This cluster had many genes encoding for innate immunity-related proteins, signaling-related, and receptors including ion channels. Cluster b also comprised several genes associated with xenobiotic metabolism. This cluster is reflective of the mantle exposure to external factors and the adductor muscle use of ion potentials for opening and closing functions (Supplementary Table 2)

A third cluster (cluster c) grouped genes that were very highly expressed in adductor tissues in contrast with brooding clutches and gonad (87 genes; 45% unknown function; 2% unannotated). Cluster c had specific muscle contracting and structural genes such as actin, twitching-like, and filamin, as well as ion channels and multiple receptors. A fourth cluster (cluster d) had 12 genes highly expressed in mantle tissues, only two of which has a functional annotation. A final cluster (cluster e) also consisted of genes with higher expression in mantle but with even more differentiation from brooding clutches (738 genes, 59.9% unknown function, 7% no annotation). Cluster e was somewhat comparable in content to cluster b, with many genes encoding products related to immunity, defense, and interaction with the environment (e.g. complement C1q tumor necrosis factor, heat shock proteins, chitinase, apoptosis-related genes). This was the only cluster to contain cytochrome P450 genes (CYP), specifically from the CYP2 and CYP3 clans, which are heavily involved in detoxification processes in invertebrates. Genes encoding for perlucin were also only found in this cluster (Figure 4B, Supplementary Table 2).

## Conclusion

Using Oxford Nanopore ultra-long read and Hi-C sequence data, the *O. lurida* genome was assembled to 10 chromosomes and a total assembly length of 1.03 Gb. The transcript diversity captured in the tissue-specific RNA-seq resulted in the annotation of 52,714 genes, 33,912 of which were fully supported which is about 16,000 and 6,000 more than those in the *O. edulis* reference genome. All 10 chromosomes of *O. lurida* were highly syntenic of all 10 chromosomes of *O. edulis* but with high levels of intra-chromosomal translocations. A single candidate structural variation region was identified by aligning ONT reads with the final assembly. Tissues showed distinct gene expression with mantle showing signatures of tissues in the interphase with external environments. Although gene prediction at the structural level was comprehensive, too much remains unknown about oysters’ and mollusk protein function. The new resource we provide here will greatly expand opportunities to increase our understanding on Olympia oysters and their interaction with the environment, as well as aid in restoration efforts and serve to uncover biological details of related bivalves.

## Data availability

The primary assembly was submitted to the National Center for Biotechnology Information (NCBI) under BioProject accession number PRJNA1504464. The individual used for WGS sequencing and assembly was assigned the Darwin Tree of Life ID xbOstLuri2 and NCBI BioSample accession number SAMN62059163. Raw reads were deposited into the NCBI Sequence Read Archive (SRA) under accessions SRR39910662 (ONT WGS), SRR39910661 (Element Hi-C), and SRR39910651-SRR39910660 (Element RNA-seq). The authors affirm that all data necessary for confirming the conclusions of the article are present within the article, figures, tables, supplementary data, or in public repositories as indicated.

## Funding

This research was supported by the U.S. Department of Agriculture, Agricultural Research Service (Project numbers: 2076-63000-005-000-D and 2040-30400-003-000-D).

This research used resources provided by the SCINet project and/or the AI Center of Excellence of the USDA Agricultural Research Service, ARS project numbers 0201-88888-003-000D and 0201-88888-002-000D.

USDA is an equal opportunity provider, employer, and lender.

## References

Astashyn, A., Tvedte, E. S., Sweeney, D., Sapojnikov, V., Bouk, N., Joukov, V., Mozes, E., Strope, P. K., Sylla, P. M., Wagner, L., Bidwell, S. L., Brown, L. C., Clark, K., Davis, E. W., Smith-White, B., Hlavina, W., Pruitt, K. D., Schneider, V. A., & Murphy, T. D. (2024). Rapid and sensitive detection of genome contamination at scale with FCS-GX. Genome Biology, 25(1), 60. 10.1186/s13059-024-03198-7

Baker, P. (1995). Review of ecology and fishery of the olympia oyster, *Ostrea lurida* with annotated bibliography. Journal of Shellfish Research, 14(2), 501–518. https://www.resalliance.org/publications/119

Barber, J. S., Greiner, C. M., & Grossman, S. (2015). Olympia oyster, Ostrea lurida, pilot project in Northern Puget Sound, Washington: 2014 monitoring report (SWIN-TR-2015-01). https://www.swinomish-nsn.gov/media/376

Bean, T. P., Tanguy, A., Peñaloza, C., Gundappa, M. K., Boutet, I., Houston, R. D., Macqueen, D. J., & Boudry, P. (2022). Two parallel chromosome-level reference genomes to support restoration and aquaculture of European flat oyster *Ostrea edulis*. Evolutionary Applications, 15(11), 1709–1712. 10.1111/eva.13465

Blake, B., & Bradbury, A. (2012). Plan for rebuilding Olympia oyster (Ostrea lurida) populations in Puget sound with a historical and contemporary overview. https://restorationfund.org/wp-content/uploads/2019/02/olympia_oyster_restoration_plan_final.pdf

Blake, B., & Ermgassen, P. S. E. Z. (2015). The history and decline of *Ostrea lurida* in Willapa Bay, Washington. Journal of Shellfish Research, 34(2), 273–280, 278. 10.2983/035.034.0208

Boutet, I., Alves Monteiro, H. J., Baudry, L., Takeuchi, T., Bonnivard, E., Billoud, B., Farhat, S., Gonzales-Araya, R., Salaun, B., Andersen, A. C., Toullec, J.-Y., Lallier, F. H., Flot, J.-F., Guiglielmoni, N., Guo, X., Li, C., Allam, B., Pales-Espinosa, E., Hemmer-Hansen, J.,…Tanguy, A. (2022). Chromosomal assembly of the flat oyster (*Ostrea edulis* L.) genome as a new genetic resource for aquaculture. Evolutionary Applications, 15(11), 1730–1748. 10.1111/eva.13462

Challis, R., Richards, E., Rajan, J., Cochrane, G., & Blaxter, M. (2020). BlobToolKit – interactive quality assessment of genome assemblies. G3 Genes|Genomes|Genetics, 10(4), 1361-1374. 10.1534/g3.119.400908

Cheng, H., Concepcion, G. T., Feng, X., Zhang, H., & Li, H. (2021). Haplotype-resolved *de novo* assembly using phased assembly graphs with hifiasm. Nature Methods, 18(2), 170–175. 10.1038/s41592-020-01056-5

Cheng, H., Jarvis, E. D., Fedrigo, O., Koepfli, K.-P., Urban, L., Gemmell, N. J., & Li, H. (2022). Haplotype-resolved assembly of diploid genomes without parental data. Nature Biotechnology, 40(9), 1332–1335. 10.1038/s41587-022-01261-x

Coe, W. R. (1932). Development of the gonads and the sequence of the sexual phases in the California oyster (Ostrea lurida). University of California Press.

Danecek, P., Bonfield, J. K., Liddle, J., Marshall, J., Ohan, V., Pollard, M. O., Whitwham, A., Keane, T., McCarthy, S. A., Davies, R. M., & Li, H. (2021). Twelve years of SAMtools and BCFtools. GigaScience, 10(2), giab008. 10.1093/gigascience/giab008

Durand, N. C., Robinson, J. T., Shamim, M. S., Machol, I., Mesirov, J. P., Lander, E. S., & Aiden, E. L. (2016). Juicebox provides a visualization system for Hi-C contact maps with unlimited zoom. Cell Systems, 3(1), 99–101. 10.1016/j.cels.2015.07.012

Echtle, E. (2014). Cultural History of the Olympia Oyster. Olympia Historical Socienty & Bigelow House Museum. https://olympiahistory.org/cultural-history-of-the-olympia-oyster-by-ed-echtle/

Emms, D. M., & Kelly, S. (2019). OrthoFinder: phylogenetic orthology inference for comparative genomics. Genome Biology, 20(1), 238. 10.1186/s13059-019-1832-y

Friedman, C. S., Brown, H. M., Ewing, T. W., Griffin, F. J., & Cherr, G. N. (2005). Pilot study of the Olympia oyster *Ostrea conchaphila* in the San Francisco Bay estuary: Description and distribution of diseases. Diseases of Aquatic Organisms, 65, 1–8. 10.3354/dao065001

Girgis, H. Z. (2015). Red: an intelligent, rapid, accurate tool for detecting repeats de-novo on the genomic scale. BMC Bioinformatics, 16(1), 227. 10.1186/s12859-015-0654-5

Gray, M., Van Acker, L., Carmele-Rescant, D., Havenhand, J., Kinnby, A., & De Wit, P. (2026). Brood chamber carbonate chemistry in the oyster *Ostrea edulis* is dynamically shaped by warming, acidification, and ventilation. Global Change Biology, 32(8), e71007. 10.1111/gcb.71007

Gray, M., W., Chaparro, O., Huebert, K., B., O’Neill, S., P., Couture, T., Moreira, A., & Brady, D. C. (2019). Life history traits conferring larval resistance against ocean acidification: The case of brooding oysters of the genus *Ostrea*. Journal of Shellfish Research, 38(3), 751–761. 10.2983/035.038.0326

Grier, C., Angelbeck, B., & McLay, E. (2017). Terraforming and monumentality as long-term social practice in the Salish Sea region of the Northwest Coast of North America. Hunter Gatherer Research, 3(1), 107–132. 10.3828/hgr.2017.7

Guan, D., McCarthy, S. A., Wood, J., Howe, K., Wang, Y., & Durbin, R. (2020). Identifying and removing haplotypic duplication in primary genome assemblies. Bioinformatics, 36(9), 2896–2898. 10.1093/bioinformatics/btaa025

Guo, X., Puritz, J. B., Wang, Z., Proestou, D., Allen, S., Small, J., Verbyla, K., Zhao, H., Haggard, J., Chriss, N., Zeng, D., Lundgren, K., Allam, B., Bushek, D., Gomez-Chiarri, M., Hare, M., Hollenbeck, C., La Peyre, J., Liu, M.,…Wilbur, A. (2023). Development and evaluation of high-density snp arrays for the Eastern oyster *Crassostrea virginica*. Marine Biotechnology, 25(1), 174–191. 10.1007/s10126-022-10191-3

Kolde, R. (2025). pheatmap: Pretty Heatmaps. In (Version 1.0.13) https://github.com/raivokolde/pheatmap

Li, H. (2018). Minimap2: pairwise alignment for nucleotide sequences. Bioinformatics, 34(18), 3094–3100. 10.1093/bioinformatics/bty191

Love, M. I., Huber, W., & Anders, S. (2014). Moderated estimation of fold change and dispersion for RNA-seq data with DESeq2. Genome Biology, 15(12), 550. 10.1186/s13059-014-0550-8

Lovell, J. T., Sreedasyam, A., Schranz, M. E., Wilson, M., Carlson, J. W., Harkess, A., Emms, D., Goodstein, D. M., & Schmutz, J. (2022). GENESPACE tracks regions of interest and gene copy number variation across multiple genomes. eLife, 11, e78526. 10.7554/eLife.78526

Luby, E. M., & Gruber, M. F. (1999). The dead must be fed: Symbolic meanings of the shellmounds of the San Francisco Bay area. Cambridge archaeological journal, 9(1), 95–108. 10.1017/S0959774300015225

Manni, M., Berkeley, M. R., Seppey, M., Simão, F. A., & Zdobnov, E. M. (2021). BUSCO Update: Novel and Streamlined Workflows along with Broader and Deeper Phylogenetic Coverage for Scoring of Eukaryotic, Prokaryotic, and Viral Genomes. Molecular Biology and Evolution, 38(10), 4647–4654. 10.1093/molbev/msab199

Miller, S. A., Dykes, D. D., & Polesky, H. F. (1988). A simple salting out procedure for extracting DNA from human nucleated cells. Nucleic Acids Research, 16(3), 1215–1215. 10.1093/nar/16.3.1215

Patro, R., Duggal, G., Love, M. I., Irizarry, R. A., & Kingsford, C. (2017). Salmon provides fast and bias-aware quantification of transcript expression. Nature Methods, 14(4), 417–419. 10.1038/nmeth.4197

Paulin, L. F., Romanek, H., Jaryani, F., & Sedlazeck, F. J. (2026). Structural variant calling using Sniffles2. Nature Protocols. 10.1038/s41596-026-01362-w

Pedersen, B. S., & Quinlan, A. R. (2018). Mosdepth: quick coverage calculation for genomes and exomes. Bioinformatics, 34(5), 867–868. 10.1093/bioinformatics/btx699

Pritchard, C., Shanks, A., Rimler, R., Oates, M., & Rumrill, S. (2015). The Olympia oyster *Ostrea lurida*: Recent advances in natural history, ecology, and restoration. Journal of Shellfish Research, 34(2), 259–271. 10.2983/035.034.0207

Puritz, J. B., Guo, X., Hare, M., He, Y., Hillier, L. W., Jin, S., Liu, M., Lotterhos, K. E., Minx, P., Modak, T., Proestou, D., Rice, E. S., Tomlinson, C., Warren, W. C., Witkop, E., Zhao, H., & Gomez-Chiarri, M. (2024). A second unveiling: Haplotig masking of the Eastern oyster genome improves population-level inference. Molecular Ecology Resources, 24(1), e13801. 10.1111/1755-0998.13801

R Core Team, R. (2026). R: A Language and Environment for Statistical Computing. In R Foundation for Statistical Computing. https://www.R-project.org/

Rausch, T., Zichner, T., Schlattl, A., Stütz, A. M., Benes, V., & Korbel, J. O. (2012). DELLY: structural variant discovery by integrated paired-end and split-read analysis. Bioinformatics, 28(18), i333–i339. 10.1093/bioinformatics/bts378

Reeder-Myers, L., Braje, T. J., Hofman, C. A., Elliott Smith, E. A., Garland, C. J., Grone, M., Hadden, C. S., Hatch, M., Hunt, T., Kelley, A., LeFebvre, M. J., Lockman, M., McKechnie, I., McNiven, I. J., Newsom, B., Pluckhahn, T., Sanchez, G., Schwadron, M., Smith, K. Y.,…Rick, T. C. (2022). Indigenous oyster fisheries persisted for millennia and should inform future management. Nature Communications, 13(1), 2383. 10.1038/s41467-022-29818-z

Ridlon, A. D., Marks, A., Zabin, C. J., Zacherl, D., Allen, B., Crooks, J., Fleener, G., Grosholz, E., Peabody, B., Toft, J., & Wasson, K. (2021). Conservation of marine foundation species: Learning from native oyster restoration from California to British Columbia. Estuaries and Coasts, 44(7), 1723–1743. 10.1007/s12237-021-00920-7

Ridlon, A. D., Wasson, K., Waters, T., Adams, J., Donatuto, J., Fleener, G., Froehlich, H., Govender, R., Kornbluth, A., Lorda, J., Peabody, B., Pinchot Iv, G., Rumrill, S. S., Tobin, E., Zabin, C. J., Zacherl, D., & Grosholz, E. D. (2021). Conservation aquaculture as a tool for imperiled marine species: Evaluation of opportunities and risks for Olympia oysters, *Ostrea lurida*. PLOS ONE, 16(6), e0252810. 10.1371/journal.pone.0252810

Silliman, K. (2019). Population structure, genetic connectivity, and adaptation in the Olympia oyster (Ostrea lurida) along the west coast of North America. Evolutionary Applications, 12(5), 923–939. 10.1111/eva.12766

Soneson, C., Love, M., & Robinson, M. (2016). Differential analyses for RNA-seq: transcript-level estimates improve gene-level inferences [version 2; peer review: 2 approved]. F1000Research, 4(1521). 10.12688/f1000research.7563.2

Steele, E. N. (1957). The rise and decline of the Olympia oyster. Fulco Publications.

Steele, E. N. (1964). The immigrant oyster (Ostrea gigas) now known as the Pacific oyster. Quick Print.

Tegenfeldt, F., Kuznetsov, D., Manni, M., Berkeley, M., Zdobnov, E. M., & Kriventseva, E. V. (2025). OrthoDB and BUSCO update: annotation of orthologs with wider sampling of genomes. Nucleic Acids Research, 53(D1), D516–D522. 10.1093/nar/gkae987

Trimble, A., C., Ruesink, J., L., & Dumbauld, B., R.. (2009). Factors preventing the recovery of a historically overexploited shellfish species, Ostrea lurida Carpenter 1864. Journal of Shellfish Research, 28(1), 97-106. 10.2983/035.028.0116

Vasimuddin, M., Misra, S., Li, H., & Aluru, S. (2019, 20-24 May 2019). Efficient architecture-aware acceleration of bwa-mem for multicore systems. 2019 IEEE International Parallel and Distributed Processing Symposium (IPDPS),

Vurture, G. W., Sedlazeck, F. J., Nattestad, M., Underwood, C. J., Fang, H., Gurtowski, J., & Schatz, M. C. (2017). GenomeScope: fast reference-free genome profiling from short reads. Bioinformatics, 33(14), 2202–2204. 10.1093/bioinformatics/btx153

Wasson, K., Gossard, D. J., Gardner, L., Hain, P. R., Zabin, C. J., Fork, S., Ridlon, A. D., Bible, J. M., Deck, A. K., & Hughes, B. B. (2020). A scientific framework for conservation aquaculture: A case study of oyster restoration in central California. Biological Conservation, 250, 108745. 10.1016/j.biocon.2020.108745

White, J. M., Buhle, E. R., Ruesink, J. L., & Trimble, A., C.. (2009). Evaluation of Olympia oyster (*Ostrea lurida* carpenter 1864) status and restoration techniques in Puget Sound, Washington, United States. Journal of Shellfish Research, 28(1), 107–112. 10.2983/035.028.0101

Zhou, C., McCarthy, S. A., & Durbin, R. (2023). YaHS: yet another Hi-C scaffolding tool. Bioinformatics, 39(1), btac808. 10.1093/bioinformatics/btac808

Zu Ermgassen, P. S. E., Spalding, M. D., Blake, B., Coen, L. D., Dumbauld, B., Geiger, S., Grabowski, J. H., Grizzle, R., Luckenbach, M., McGraw, K., Rodney, W., Ruesink, J. L., Powers, S. P., & Brumbaugh, R. (2012). Historical ecology with real numbers: past and present extent and biomass of an imperilled estuarine habitat. Proceedings of the Royal Society B: Biological Sciences, 279(1742), 3393–3400. 10.1098/rspb.2012.0313

